# Integration of microprism and microelectrode arrays enables simultaneous electrophysiology and two-photon imaging across all cortical layers

**DOI:** 10.1101/2022.07.08.499369

**Authors:** Qianru Yang, Bingchen Wu, Elisa Castagnola, May Yoon Pwint, Nathaniel Williams, Alberto L. Vazquez, X. Tracy Cui

**Affiliations:** Department of Bioengineering, University of Pittsburgh, United States; Center for Neural Basis of Cognition, University of Pittsburgh and Carnegie Mellon University, United States; Department of Radiology, University of Pittsburgh, United States; McGowan Institute for Regenerative Medicine, University of Pittsburgh, United States

**Keywords:** Microprism, electrophysiology, calcium imaging, microstimulation, two-photon microscopy

## Abstract

Electrophysiology is a vital tool in neuroscience research with increasing translational value. It is used to record or modulate neuronal activity with high temporal but lower spatial resolution. Optical technologies, such as two-photon microscopy (TPM) can complement electrophysiological recordings with large-scale imaging at cellular resolution. Combining these two provides a powerful platform to elucidate and coordinate multimodal functions. Prior attempts have been limited to the superficial brain from a top-down optical view. Here, we describe a novel combination of transparent microelectrode arrays (MEAs) with glass microprisms for simultaneous electrophysiology and optical imaging of all cortical layers in a vertical plane. We tested our device in Thy1-GCaMP6 mice for over 4 months and demonstrated its capability for multisite single-unit recording, microstimulation, and simultaneous TPM calcium imaging. Using this setup, we reveal how amplitude, frequency, and depth of microstimulation impact neural activation patterns across the cortical column. This work presents a multimodal tool that extends integrated electrophysiology and optical imaging from the superficial brain to the whole cortical column, opening new avenues of neuroscience research and neurotechnology development.

## 1. Introduction

Understanding complex neural networks in the brain is one of the biggest challenges in neuroscience. Electrophysiology has been a crucial method since the discovery of action potentials in the late 19^th^ century [1], which directly links neural functions with electrical activity. Microelectrode arrays (MEAs), including the Utah array and Michigan array, are the most widely used devices to record or modulate neural activity [2–9]. Studies using in vivo electrophysiology have revealed the functions of different brain regions, building the foundation for basic and translational neuroscience. However, little is known about the true spatial distribution of neurons interfacing with MEAs in vivo [10]. In addition, it is still a challenge to distinguish large numbers of individual neurons from recording data. Recent breakthroughs in optical imaging technologies, such as in vivo two-photon microscopy (TPM) [11–13], can provide important complementary information. Large-scale calcium imaging has shown dynamic activity patterns from imaging hundreds of neurons with cellular resolution [14–17]. In addition, development of endogenous or exogenous fluorescent labeling for specific molecules and/or cell types further allow analyzing a rich diversity of cellular and subcellular activity [18–25]. Combining electrophysiology with optical methods can leverage the advantages of both methodologies. For example, simultaneous electrophysiological recording and TPM imaging of different types of neurons can help us understand the contributions of neural subtypes to electrophysiology features. Simultaneous microstimulation and TPM imaging would allow us to investigate the mechanism of microstimulation on the neural circuit and population level.

There have been several attempts at simultaneous electrophysiology and optical imaging. Rigid silicon electrodes have been inserted into the brain at an angle (∼30 degrees relative to the brain surface) to enable imaging of the brain-electrode interface [26–31]. Such configurations suffer from acute bleeding caused by the implanted electrode and chronic fibrous tissue growth at the brain surface, which significantly decreases the imaging depth of TPM from ∼600 μm (no implant) to ∼ 300 μm (acute implant) or ∼ 150 μm (chronic implant) [32, 33]. The opaque electrode also blocks the light for imaging tissue underneath. Transparent electrodes have recently drawn a lot of interest due to their compatibility with optical modalities. Typically, a transparent electrode is microfabricated by patterning conductive materials on a transparent and flexible insulating substrate (e.g., parylene C, SU-8, and polyimide). Two categories of conductive materials have been utilized in electrode fabrication: transparent conductive materials such as indium tin oxide (ITO) and graphene, and non-transparent metal materials such as gold and platinum. Almost totally transparent ITO or graphene MEAs have achieved low crosstalk electrocorticography (ECoG) recording and stimulation with concurrent TPM imaging or optogenetic modulations [34–40]. However, compared to metal electrodes, the high electrochemical impedance of these transparent materials limits the miniaturization of the electrodes for recording single neuron activity. In addition, the electrical stimulation capability and stability of these materials have not yet been extensively investigated. Metal materials have long been used in interfacing with neural tissue. Platinum (Pt) and Iridium Oxide are the two “gold-standard” materials for stimulating electrodes due to their chemical inertness and high charge-storage capacity. In this work we choose Pt as our conductive material for its well-established capability of microstimulation. Several groups have explored the possibility of metal-based transparent MEAs [41–43]. Due to the non-transparent property of metal materials, this kind of MEA seeks to minimize the metal traces while adding low impedance coatings to improve their electrochemical performance. For example, Qiang et al. designed a transparent gold (Au) nano-mesh MEA with a poly(3,4-ethylene dioxythiophene)-poly(styrene sulfonate) (PEDOT: PSS) coating. This metal-based transparent MEA outperformed previous graphene and ITO microelectrodes in both impedance and charge injection limit (CIL) by more than one order of magnitude, at the cost of reduced optical transparency (67%) [43]. Despite these achievements, all previous work has been optically limited to the superficial brain (layers 1-3). Although the laminar structure of the cortex has long been known and is widely used in building computational neural networks [44–47], the dynamics of cortical microcircuits among large numbers of neurons across layers remain poorly understood [48]. A methodology to monitor and modulate neuron populations in both superficial and deep layers simultaneously in vivo could greatly promote our understanding of the cortical neural microcircuit across laminar depth.

Here we describe an approach that integrates a transparent MEA with a glass microprism, enabling electrophysiology and simultaneous optical imaging from a vertical view throughout all cortical layers. The transparent MEA achieved chronic in vivo electrophysiology recording and microstimulation by using platinum (Pt) as the conductive material combined with a nano Pt coating. The transparency of the MEA is achieved with a transparent SU-8 substrate and thin metal traces. We adopted a ring shape for the electrode sites to minimize the optical shadow. This novel design provided a facile way to manufacture mostly transparent MEAs with non-transparent conducting materials. Whole cortical column imaging is realized by the microprism implantation, which changes the light path by 90 degrees to present a vertical view [49–52]. Although there are concerns about tissue damage caused by implanting a 1 mm microprism, our group has previously shown using TPM over a 32 week chronic implantation period that the microglia response to microprism implants subsides and brain tissue returns to normal within 4-8 weeks [53]. In this work, we developed a process to assemble a functional transparent MEA onto the microprism, and then implant the integrated device as a whole to avoid the additional trauma of electrode insertion. We tested this MEA-on-microprism device in Thy1-GCaMP6 mice for over 4 months, performing intracortical single unit neural recording, microstimulation, and large-scale two-photon calcium imaging from the brain surface to cortical layer 5/6. In addition, we investigated how parameters and depth of microstimulation affect neural population activity using this setup. Through simultaneous microstimulation and TPM imaging, we acquired the first suite of calcium images from neuronal populations across multiple cortical layers in response to microstimulation testing under a series of parameters, providing new insights for choosing microstimulation parameters and depth in neuroscience research and neural prosthetics applications.

## 2. Results and Discussions

To enable the integration of MEAs with the microprism, we custom-designed and fabricated transparent MEAs using standard photolithography techniques (details in Materials and Methods and Fig. 1a-e). The 16-channel MEA used SU-8 as the insulating substrate and Pt as the conductive material. SU-8 has been extensively utilized in biomedical devices due to its high transparency, low young’s modulus, and high yield strength [54–56]. Pt is a common material for stimulation electrodes for its chemical inertness [57–59]. The insulating SU-8 layers were optimized to be 9 µm thick on each side, balancing between flexibility and robustness. To allow optical imaging through the MEA, the width of the metal connections was minimized to 2 µm within the 1 x 1 mm2 tip region attached to the imaging face of the microprism, and for the electrode sites we adopted a ring-shaped design (Fig. 1f, g). The ring-shaped sites had an external diameter of 50 µm and an internal diameter of 40 µm, to match the geometric area (706 µm2) with a standard 30 µm-diameter disk electrode. Compared to disk sites with the same geometric area, ring sites not only dramatically reduced the shadow (Fig. 1h), resolving more details of fine structures (Fig. 1i), but also showed a significantly decreased photo-electric artifact (Fig. 1j)). The 16 channels were arranged in a 4×4 configuration with an inter-electrode spacing of 200 µm (center-to-center) (Fig. 1g). The flexibility and mechanical durability of the MEA was validated with bending tests. To conduct the bending tests, the part of the MEA containing the traces was pushed against a rectangular corner with a thin metal wire (d=0.18mm) to create a 90-degree bend (Fig. 1k). After 100 repetitions of bending, the CV curve and the impedance spectrum of all three MEAs tested remained almost identical to measurements taken prior to bending (Fig. 1l, m), demonstrating that these MEAs can well survive the handling and bending needed to assemble the device.

**Figure 1.**
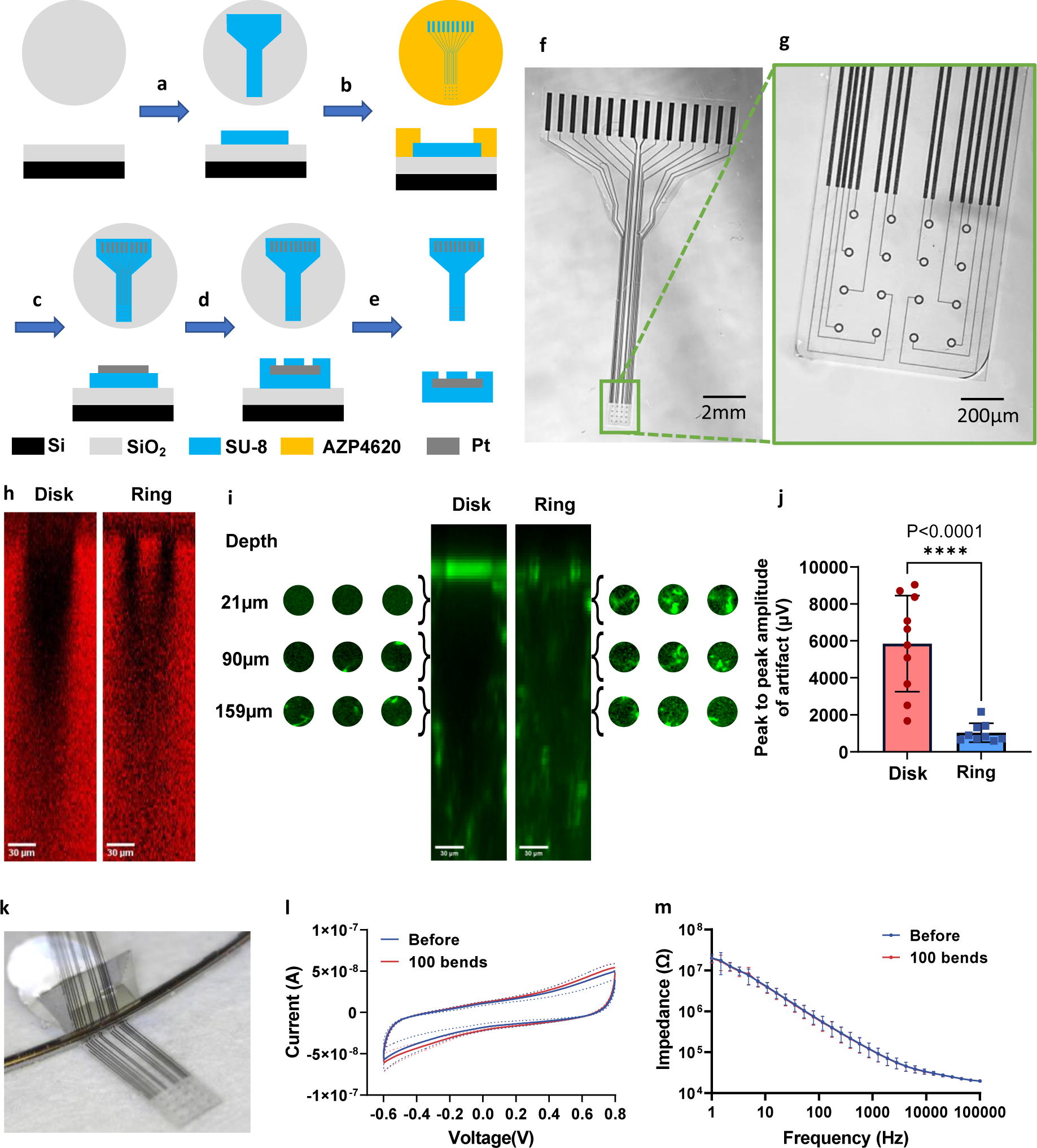
Fabrication and tests of the flexible and transparent MEA. (a)-(e) Schematic of photolithography procedures. The top and side views of each step are shown in stacks (dimensions not to scale). (a) SU-8 was spin coated, exposed, and then developed as the bottom insulation on a silicon (Si) wafer with 1µm of silicon dioxide (SiO_2_) on the surface. (b) AZ4620 positive photoresist was spin-coated, exposed, and developed as the mask for the metal layer. (c) 100 nm Platinum (Pt) was evaporated on the wafer with a 10 nm Ti adhesion layer and lifted off in acetone to form the conductive metal part. (d) Another layer of SU-8 was spin coated, exposed, and developed as the top insulation. (e) The wafer was placed in buffered oxide etchant to release the MEAs. (f) Optical microscopy image of the whole MEA. (g) Zoomed-in image of the MEA tip showing the 4×4 ring-shaped array and the thin connecting traces. (h) Representative orthogonal view of a disk (left) and ring (right) site in an SR101 gel phantom, showing the relative shadow cast by each through a uniform background. (i) Orthogonal view of the average projection for 9 disk sites (left) and 9 ring sites (right) in a CX3CR1-GFP mouse. Slices at 3 different depths are shown for 3 representative electrode sites for disk and ring arrays, showing the difference in fine process resolution for microglia under the electrode sites. (j) Peak to peak amplitude of photo-electric artifact recorded by disk sites and ring sites under the same two-photon laser scan in vitro. (k) A photograph of the bending test. The MEA was bent 90 degrees by pushing a metal wire against a rectangular corner. (l) CV curve before and after 100 bends at 90 degrees. N=31 sites from 3 MEAs. Data are presented as mean ± SD. (m) Impedance spectrum before and after 100 bends at 90 degrees. N=31 sites from 3 MEAs. Data are presented as mean ± SD.

We electrochemically deposited nano-Pt coatings on the MEAs to improve their electrochemical properties (Fig. 2a). The magnified SEM of the nano-Pt coating showed a cauliflower-like nanostructured morphology, while the surface of the bare Pt electrode was relatively flat. These images clearly show that the nano-Pt coating drastically increases the effective electrochemical area of the electrode. Consistent with the SEM results, we observed a significant increase in the charge storage capacity (CSC) of the electrode from 1.06±0.15 mC cm^−2^ to 6.05±0.17 mC cm^−2^, as calculated from the time integral of the last cyclic voltammogram (CV) cycle before and after the nano-Pt coating (Fig. 2b). We also observed that the nano-Pt coating decreased the impedance of the electrode in the low-frequency range (Fig. 2c). Specifically, the impedance at 1kHz dropped from 75.32±23.70 kΩ to 41.34±2.96 kΩ. After nano-Pt deposition, the CIL of the electrode was estimated to be 2.27 mC cm^−2^, as we could deliver 80 µA cathodic current for 200 µs before the E_mc_ reached −0.6V versus Ag/AgCl (Fig. 2d). Compared with other state-of-the-art transparent electrode arrays, our nano-Pt coated ring electrodes showed superb impedance, slightly outperformed the Au/PEDOT:PSS nanomesh microelectrodes and were 30 to 100 times better than graphene or ITO electrodes scaled to the same size [35, 36, 40, 43, 60] (Fig. 2e, supplementary table 1).

**Figure 2.**
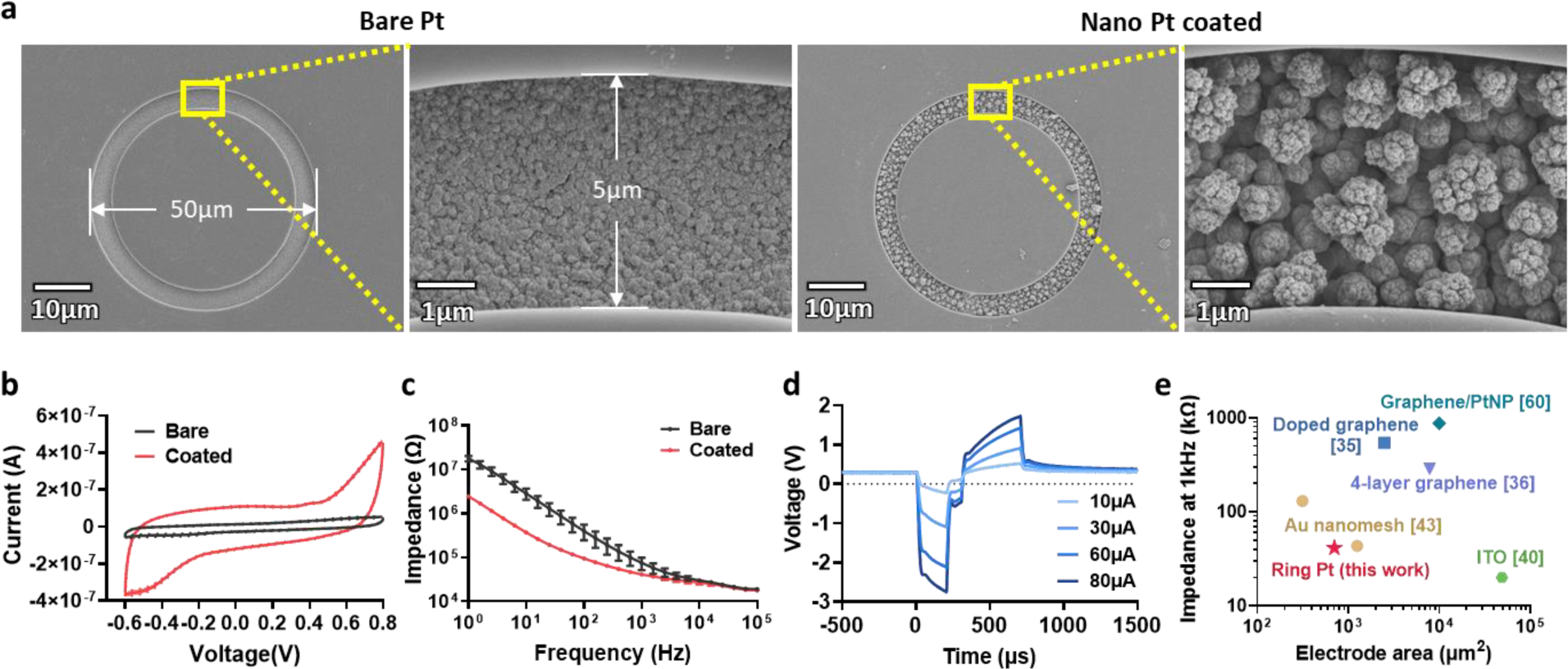
In vitro characterization of MEA before and after nano-Pt deposition. (a) Representative SEM images of bare Pt and nano-Pt coated electrode sites. Magnified views on the right side in each condition show the nanostructured surface morphologies. (b) CV of a representative MEA before and after nano-Pt coating. N=16 electrode sites. Data are presented as mean ± SD. (c) Electrochemical impedance spectroscopy (EIS) of an MEA before and after nano Pt coating. N=16 electrode sites. Data are presented as mean ± SD. (d) Voltage transients in response to bi-phasic, charge-balanced, asymmetric (I_c_= 2 * I_a_) current pulses. The cathodic current amplitudes were ramped up from 10 µA to 80 µA. (e) The impedance at 1kHz and electrode area of our MEAs compared with other state-of-art transparent MEAs.

The procedure to assemble a functional microprism/MEA device is illustrated in Fig. 3. We designed the 16-channel PCB that connects the Omnetics connector with 16 metal pads and two metal rings for the ground/reference electrode. The MEA was bonded to the PCB with an anisotropic conductive film (ACF) cable. We tested the CV and EIS of the MEA to ensure proper connections and electrochemically deposited nano-Pt onto the Pt electrode sites. Meanwhile, the microprism was attached to three pieces of coverglass by UV curable transparent optical glue. We then adhered the tip of the MEA to the microprism and the cover glass, also with optical glue. Note that the nano-Pt coating needed to be done before we glued the MEA to the microprism, otherwise the aluminum coating on the microprism might be dissolved away in the H_2_PtCl_6_ coating solution. Finally, we soldered a silver wire to the ground and reference pads on the PCB and applied a thin layer of silicone onto the traces and wires to strengthen the insulation and the mechanical durability of the device. Before the implantation surgery, we ran electrochemical characterizations of the fully assembled device and sterilized it with ethylene oxide sterilization.

**Figure 3.**
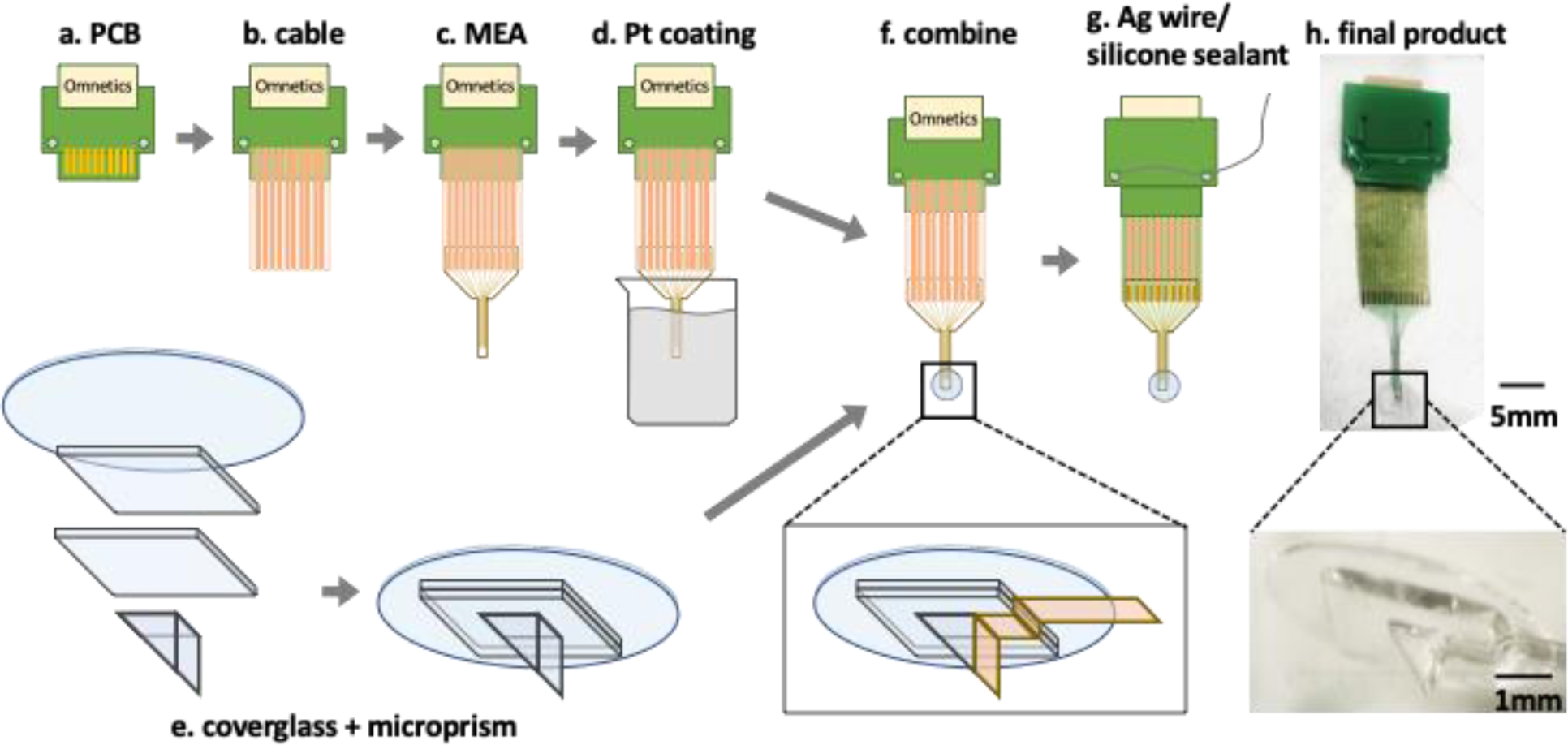
Assembly procedure of the functional MEA-on-microprism device. (a) Customized PCB with standard 16-channel Omnetics connector. (b) Connected the heat-seal ACF cable to the metal traces on the PCB. (c) Bonded our previously fabricated MEA to the ACF cable. (d) Electrochemically deposited nano-Pt onto the MEA by applying a constant voltage. (e) Glued microprism and three pieces of cover glass together using UV-cured transparent glue. (f) Adhered the tip of the MEA to the microprism imaging face using transparent glue. (g) Soldered a silver wire to the PCB as the ground/reference. Spread a thin layer of silicone sealant onto the backside of the cable, the trace region on the MEA, and covered all the exposed bare metal parts. (h) Photo of a finished assembly. The inset reveals the magnified view of the MEA tip on the microprism. Scale bar=5 mm for the top photo, and 1mm for the inset.

To test the capability of our device in vivo, we implanted the MEA-on-microprism in the visual cortex of mice for over 4 months. The surgical procedure was similar to implanting just the microprism as described in previous publications [49, 50, 53]. A metal head frame was attached to the skull for head-fixed experiments later. We created a 4mm x 4mm square cranial window in the right hemisphere between sutures in GCaMP6s mice, generally over the sensory and visual cortex of the animal. The orientation of the microprism was pre-designed to target the visual cortex during assembly. After removing dura matter with fine-tip tweezers and a bent needle (30 G), we made a 1mm long and 1mm deep incision with a reshaped razor blade (Electron Microscopy Sciences). The microprism was held with a vacuum line which was bound to a stereotaxic arm and slid vertically into the cut. Another stereotaxic arm that held the PCB approached the brain at an angle (∼30 degrees) at a synchronized pace with the vacuum arm. After inserting the device fully into the brain, we sealed the cranial window with silicone (Kwik-sil) and dental cement in sequence on the edges. The MEA was grounded to a stainless-steel bone screw on the skull on the contralateral side. Extra ACF cable was folded under PCB and the device outside of the brain was fixed to the skull by embedding in dental cement (Fig. 4a).

**Figure 4.**
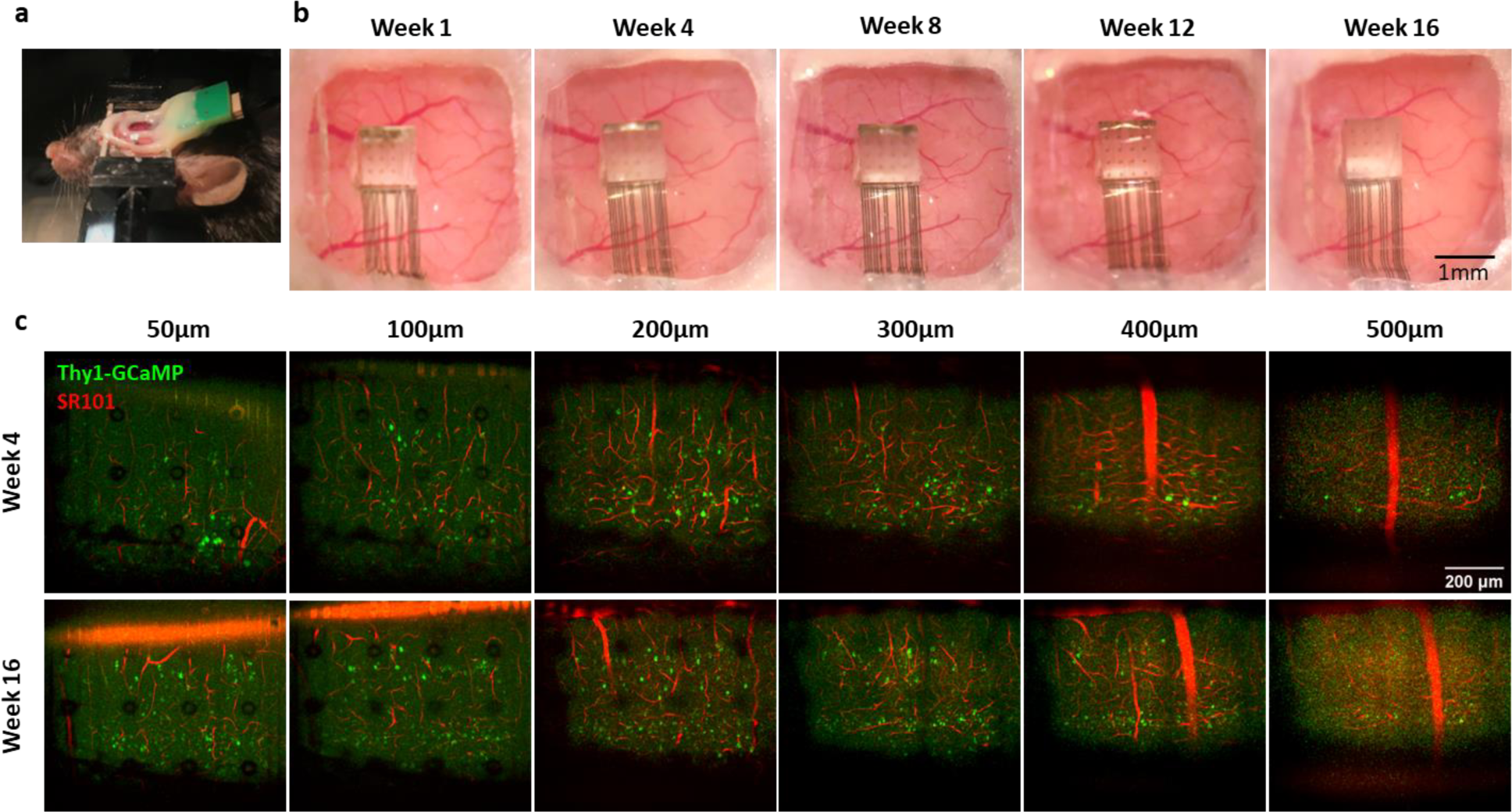
The MEA-on-microprism device implanted in mice allows high quality optical imaging over 16 weeks. (a) Bird’s eye view of the cranial window and the device on a head-fixed mouse. (b) Images of the cranial window under a surgical microscope for over 16 weeks. (c) TPM imaging of spontaneous neural activity (green) and blood vessels (red) through the MEA-on-microprism device. Neural calcium activity image is a max projection from a 3 min. time-lapse scan. Left to right: increasing distances from the microprism.

Successful surgery can maintain a clear and stable cranial window for a long time (Fig. 4b). Through the implanted microprism, we were able to monitor neuronal activity from the brain surface down to layer 5/6, confirmed by the void in the expression of layer 4 neurons in Thy1-GCaMP6s animals (Fig. 4c). Aligned with previous work [53], the imaging distance was around 500 µm from the microprism vertical face after several weeks of recovery. The ring-shaped design of the MEA reduced the width of the metal to the scale of micro blood vessels, minimizing the shadow of the metal traces while keeping the same surface area as the standard 30 µm disc electrode sites (Fig. 4c). In TPM images, the shadow of the thin metal connection is almost invisible beyond 50 µm from the microprism and the shadow of the electrode sites becomes invisible around 200 µm away from the microprism (Fig. 4c).

We characterized the electrochemical properties of the MEAs implanted in three animals using a two-electrode setup, where a stainless-steel bone screw on the contralateral brain hemisphere served as the counter electrode. Note that for in vitro electrochemical measurements we used a three-electrodes setup, thus the results are not directly comparable. We analyzed 22 electrode sites from three animals that maintained a reasonable 1kHz impedance (<1 MΩ) for the majority of the electrophysiology study. The CSC of individual electrode sites fluctuated over time as shown in Fig. 5a. The average CSC also possessed ups and downs over time—it increased in the first week then dropped back and reached the lowest value at week 2, followed by a slightly increasing trend afterward (Fig. 5a). The CSC at week 12 and week 16 were significantly higher than day 0 (immediately after the implantation surgery) (Fig. 5a). The mean impedance at 1 kHz of the MEAs increased in the first 2 weeks, then declined from week 2 to week 8, and increased back from week 8 to week 16 (Fig. 5b). The impedance at week 2, week 6, and week 8 were significantly different from day 0 (Fig. 5b). The shape of the CV curve and the impedance spectrum from all working electrode sites (impedance at 1 kHz<1 MΩ) in one animal are plotted over time in Figures 5c and 5d. In general, the CV curve and impedance spectrum remained stable throughout 16 weeks, showing the long-term stability of our custom-fabricated MEAs. The percentage of working channels (impedance at 1 kHz<1 MΩ) varied in different animals (Fig. 5e). The changes over time may have resulted from loose connections, tissue response, material degradation, mechanical damage, etc. On average, there were ∼50-70% of channels working.

**Figure 5.**
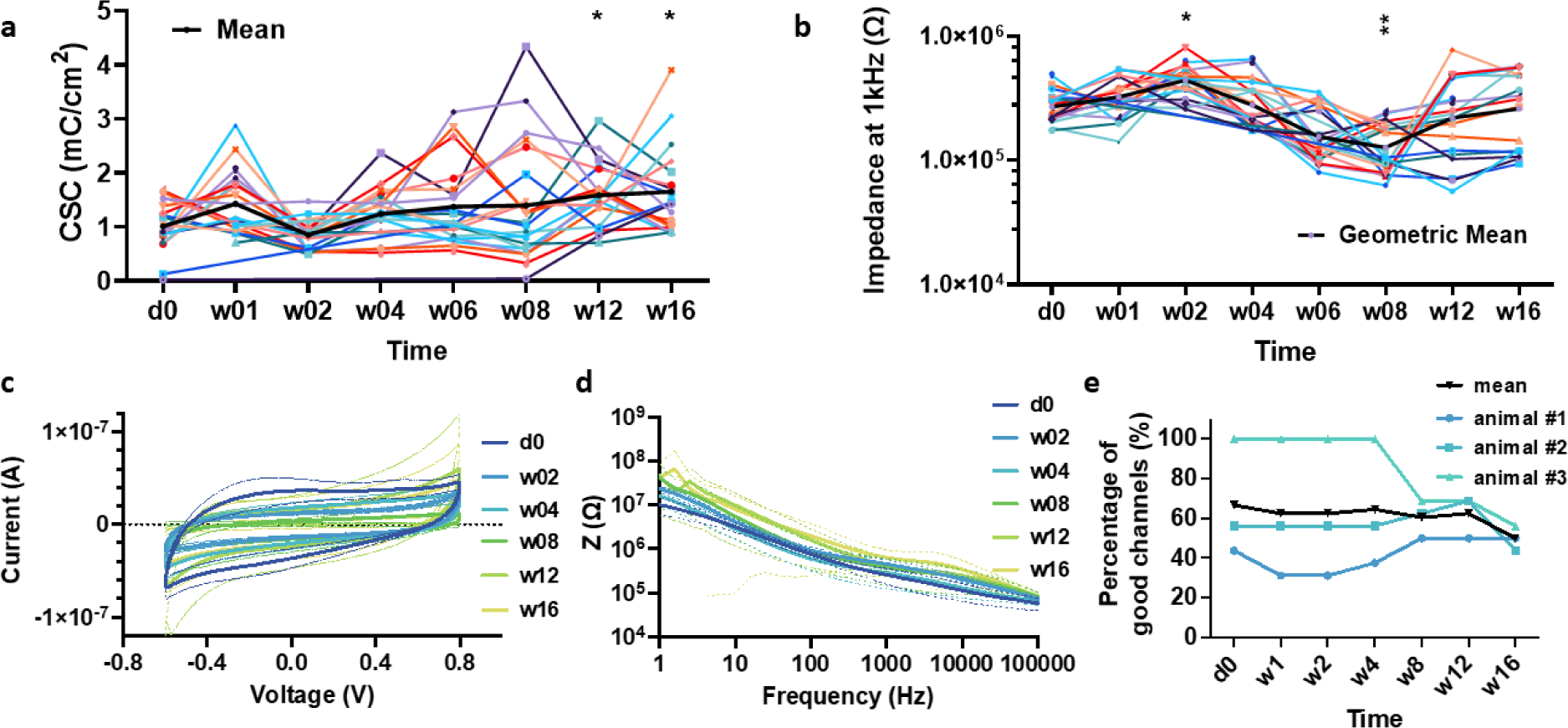
In vivo electrochemical characterization of implanted MEAs for over 16 weeks. (a) Charge storage capacity of the MEA over time (n=22 sites from 3 animals). Individual sites are plotted in assorted colors, and the mean value is presented in a thick black line. (b) The impedance at 1 kHz over time (n=22 sites from 3 animals). Individual sites are plotted in assorted colors, and the mean value is presented in a thick black line. In (a) & (b), we ran a one-way ANOVA followed by Dunn’s multiple comparisons to test for differences against time point d0. * p<0.05, ** p<0.01. (c) CV curve of all working sites (impedance at 1 kHz<1 MΩ) in one implanted MEA over time. Different colors indicate different time points. Data are presented as mean ± SD. (d) Impedance spectrum of all working sites (impedance at 1 kHz<1 MΩ) in one implanted MEA over time. Different colors indicate different time points as in (c). Data are presented as mean ± SD. (e) Summary of the percentage of working sites (impedance at 1kHz<1MΩ) in three implanted MEAs over time.

To test the electrophysiological recording capability of the MEAs, we recorded spontaneous electrophysiological activity in three awake mice for over 12 weeks. Spike train data was automatically detected by Ripple and sorted offline later. Spike units were sorted manually in 3-D PCA space using the Plexon Offline Sorter. The implanted MEAs were able to record single-unit activity in awake animals for over 12 weeks (Fig. 6a). We inspected the shape of waveforms and ensured the inter-spike intervals were larger than 1ms (Fig. 6b, c). Noise waveforms that co-occurred in multiple channels were excluded.

**Figure 6.**
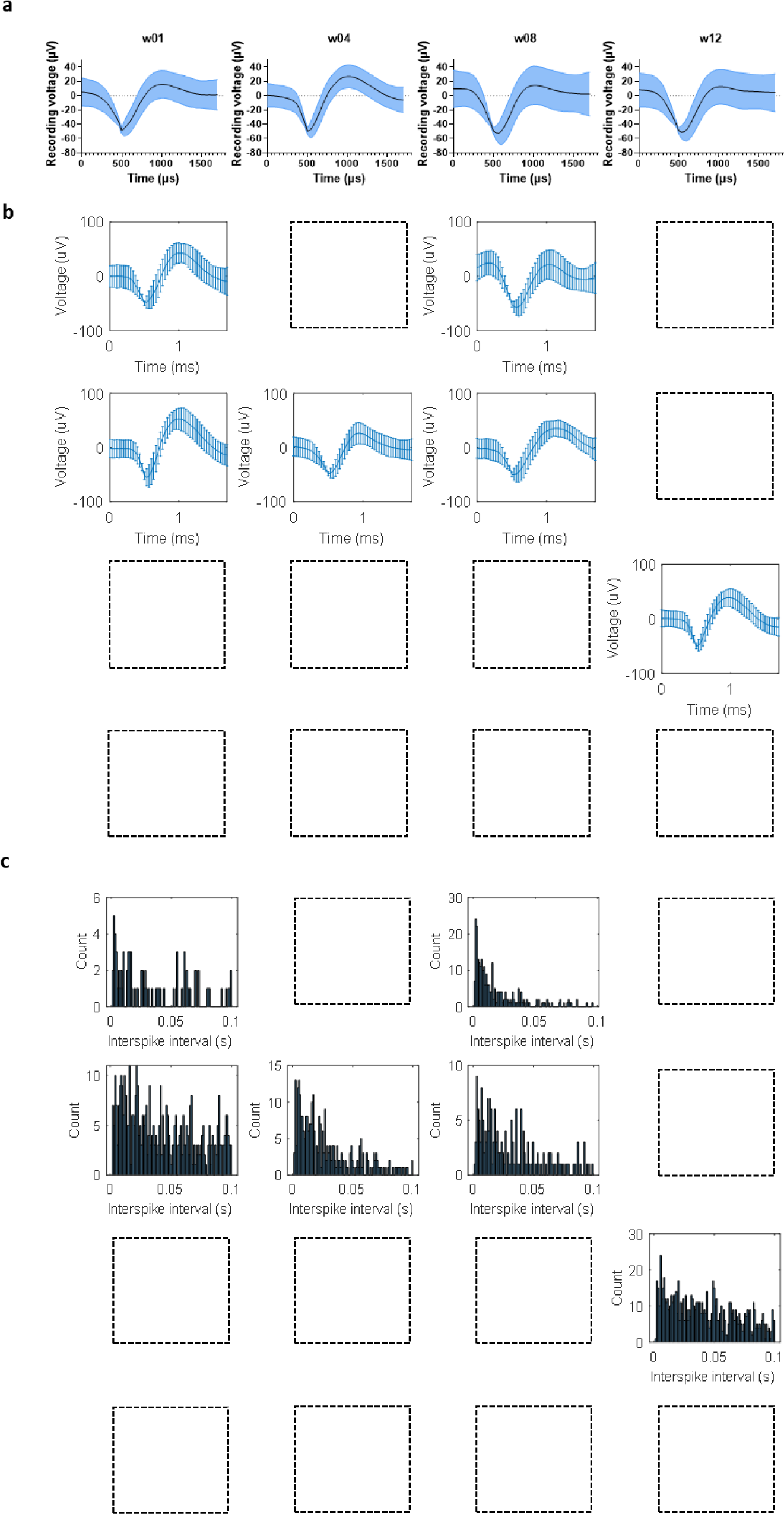
Chronic in vivo recording through the MEA. (a) Representative waveforms extracted from spontaneous recordings in an awake mouse over time. Spikes were sorted using Plexon Offline Sorter software. Data are presented as mean ± SD. (b) Spatial map of detected units in one animal at week 4 post implantation. Empty blocks indicate bad channels (impedance at 1kHz>1MΩ). The top row is superficial relative to the cortical surface. (c) Spatial map of the inter-spike intervals of detected units in (b). Empty blocks indicate bad channels (impedance at 1kHz>1MΩ).

Simultaneous recordings of spontaneous electrophysiological activity and TPM calcium fluorescence at 50µm from the MEA were also conducted with our setup. One of the major challenges of electrophysiological recording during TPM imaging is the photo-electric artifact generated by the imaging laser on the recording signal [61]. This artifact is most prominent when the laser scans over the electrode traces, such that sites that are not in the imaging plane show reduced artifact magnitude. Both negative and positive deflections were observed (Fig. 7a). Thanks to the thin traces and the ring electrode shape, the artifact is not significant enough to saturate the amplifier so that post-processing of data can proceed. The onset of each imaging frame was used to generate templates of the artifact for each channel. Features from these templates (e.g., average) were used to remove the artifact from each channel to obtain corrected recording traces (Fig. 7a). The corrected recording was then high pass filtered to extract multi-unit activity (MUA) (Fig. 7b). We compared the MUA with calcium activity and observed agreement between these traces, especially after convolving the MUA with the GCaMP6f fluorescent response function [61]. We show an example of the calcium activity from a single neuron nearby the recording electrode (Fig. 7b). Here, the correlation between the MUA and GCaMP activity was calculated to be 0.41. We performed single unit analysis of the traces recorded during TPM imaging after the artifact removal process. The same unit before and during TPM imaging showed similar ISI distributions and similar spike waveforms with slightly enlarged amplitude (Fig. 7c), indicating that this method well preserves the neural activity signal. In conclusion, this setup could be used to investigate the correlation of other cellular dynamics to electrophysiological recordings in vivo. Our current TPM was limited to single plane imaging at this resolution. Future studies that are capable of volumetric cellular imaging would be valuable to capture all cellular activity and untangle their contributions to electrophysiology signals.

**Figure 7.**
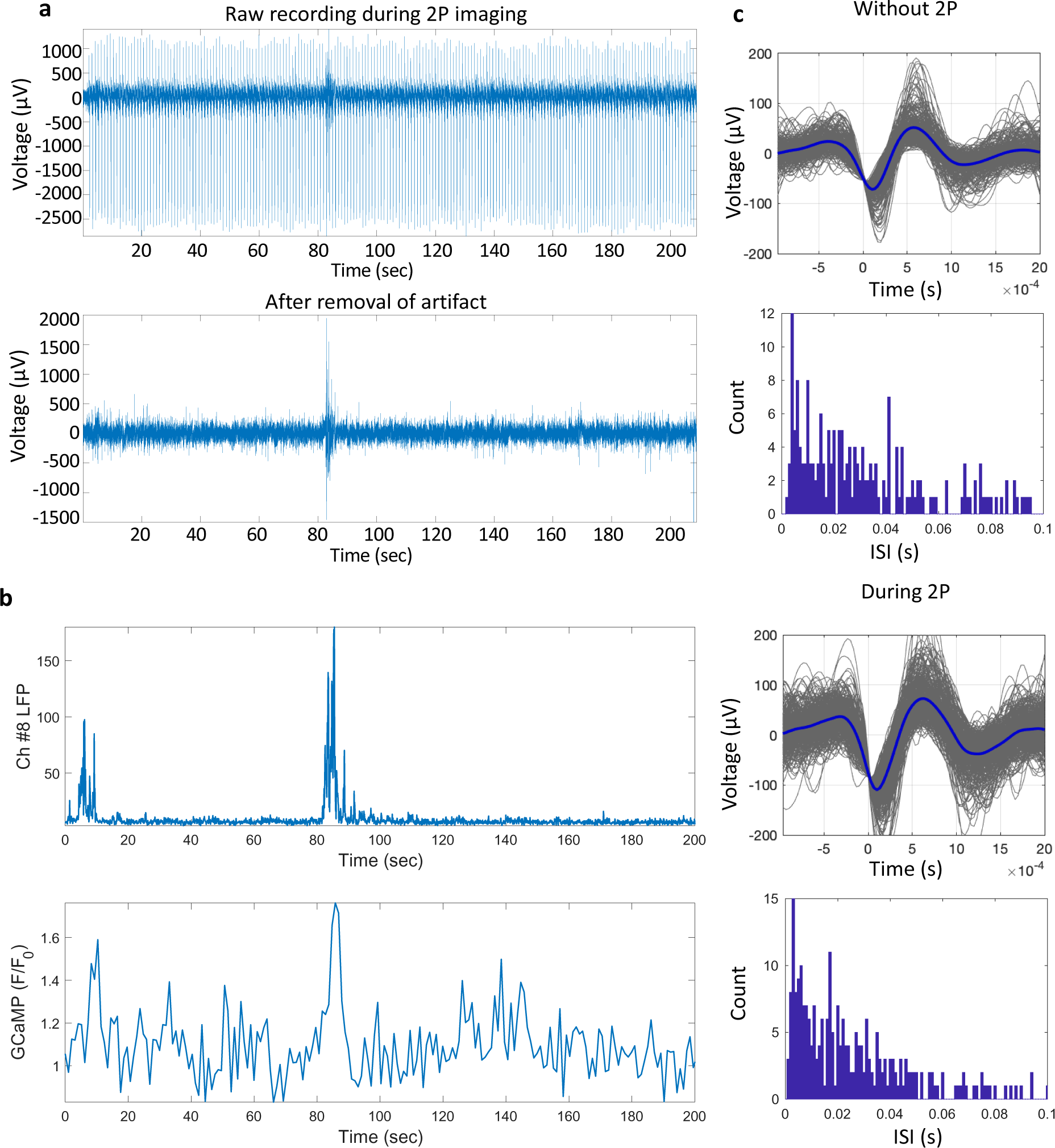
Simultaneous electrophysiology recording and TPM imaging. (a) Raw electrophysiological recording and the same recording after removal of the photo-electric artifact generated by TPM imaging. Note that the recording does not saturate the digitizer. (b) Simultaneous multi-unit activity (MUA) recorded from one channel and the TPM calcium activity of a nearby neuron. (c) The waveforms and ISI plot of a sorted unit from spontaneous recordings without two-photon and during two-photon imaging.

Next, we examined TPM imaging of neuronal activity evoked by visual stimulation to show the potential of our setup for dissecting neural circuits. A blue LED was placed about 5 cm away from the contralateral eye of the mouse. After a 30-sec delay, the LED flashed at 10 Hz for 1 second followed by a 15 second delay. By averaging over 10 trials, we identified the neurons that were repeatedly activated by the visual stimuli at 50µm from the microprism/MEA (Fig. 8a). The relative fluorescence intensity change (ΔF/F) of three representative neurons during the whole session is shown in Fig. 8b. The background fluorescence intensity (F) is the mean value of the first 30 sec before light stimulation. After aligning the curves to the time stamp of stimulus onset from the 10 trials, we found the average response of each neuron (Fig. 8c). Neuron #1 and neuron #2 peaked at the light-on period, while neuron #3 showed the largest response after light-off (Fig. 8c). These are typical responses of neurons that are sensitive to a light spot being turned on or off in the visual system [62–64]. The suppressed activity during light stimuli of neuron#3 is typical of an inhibitory intracortical circuit [65–67]. These results indicate that visual input circuit activity is preserved near the imaging face of our microprism/MEA device, showing the potential for studying a variety of sensory evoked neuronal responses with this integrated system.

**Figure 8.**
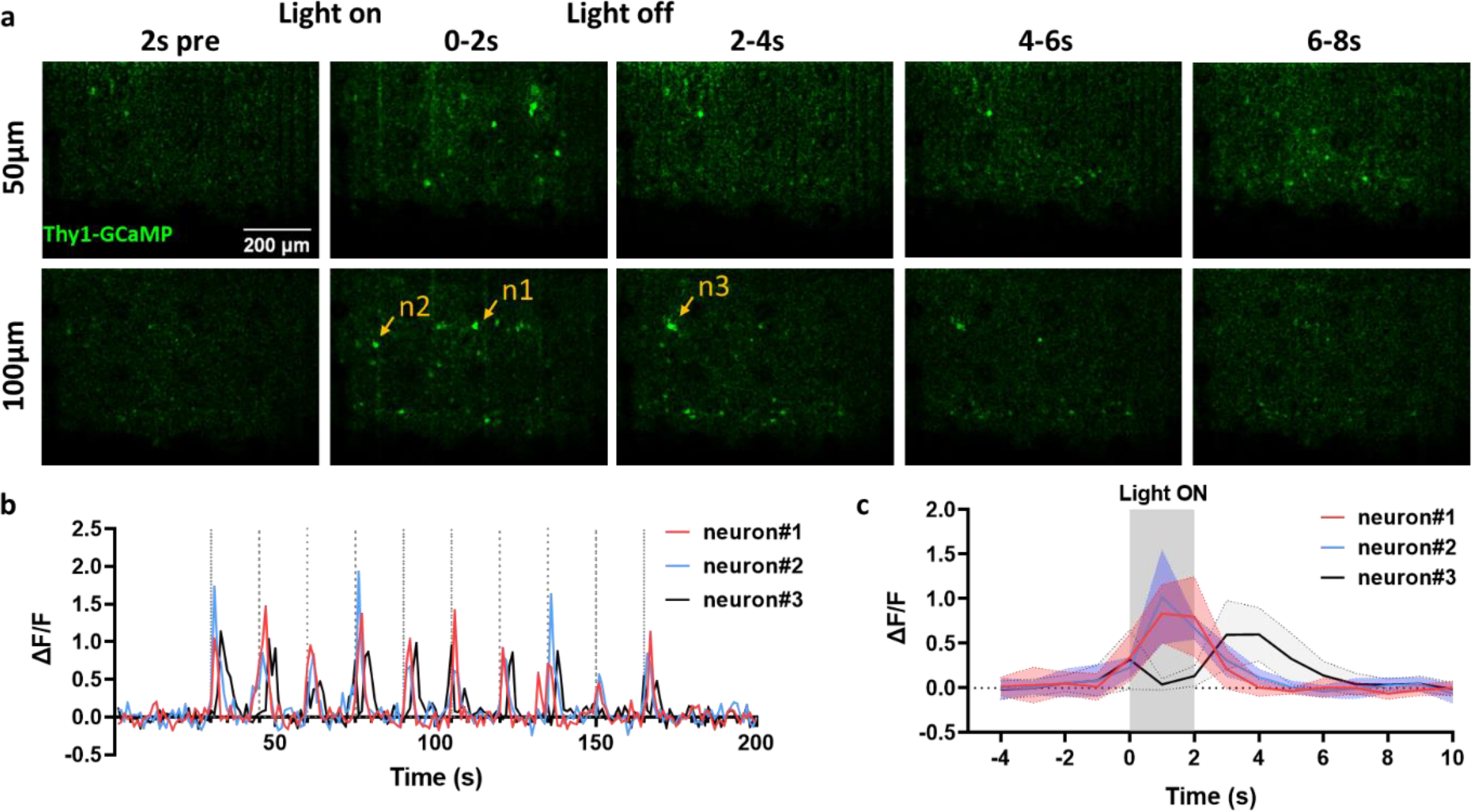
TPM imaging of visual stimulation evoked neuronal calcium response. (a) GCaMP fluorescence changes before/during/after light stimulation. Averaged from 10 consecutive trials. n1-n3 mark the three neurons analyzed in (b) and (c). (b) GCaMP fluorescence change of three representative neurons during the whole 200s imaging session. Dotted lines indicate the onset time of each light stimulation period (1 sec). (c) Average fluorescence changes of the three neurons relative to the light stimuli. Shaded ribbons represent mean ± SD.

Intracortical microstimulation has been used to restore the function of the brain and to causally study the functions of brain regions [6, 9, 57, 68]. However, the cellular mechanisms of microstimulation are not yet fully understood. For example, the identity of neurons modulated by microstimulation at different amplitudes and frequencies simultaneously across cortical layers has not been examined in vivo. Previous electrophysiological methods typically only sample a small number of neurons at a time, in addition to the technical difficulties of differentiating neural activity from the artifact of injected current. Optical imaging techniques such as TPM enable monitoring neural population activity completely free of the stimulation artifact. However, conventional TPM studies have focused on cortical layer 2/3 due to imaging depth limitations [28, 30, 69]. Using our MEA on microprism device, we recorded neuronal responses by TPM imaging throughout all cortical layers simultaneously during intracortical microstimulation. Electrical stimulation consisted of charge-balanced square wave pulses with a 200 µs cathodic phase, 100 µs interval, and 400 µs anodic phase (Fig. 9a), mimicking the parameters used in human studies [5]. The reported amplitudes here all refer to the cathodic current amplitudes. We first asked how the current amplitude of stimulation influenced neuronal activation. We stimulated every available site with an impedance of < 1MOhm at 1 kHz three times with a 25% duty cycle (1 s ON, 3 s OFF) at 50 Hz and monitored the neuronal calcium activity 50 µm away from the MEA surface using TPM in awake mice. The images were averaged from three stimulation periods. Near the activation threshold current (5 µA), a couple of sparsely distributed neurons were activated (Fig. 9b). As we increased the current amplitude to 10µA, more neurons near the stimulated electrode became activated (Fig. 9b). When the current amplitude reached 15 µA, we observed substantial neuropil activation around the stimulated site and that the neural response appeared to saturate at >15 µA (Fig. 9b). Similar neuronal calcium responses to changing amplitudes were observed on all working electrode sites over different days. The sparse and distributed neuronal activation at 5 µA is consistent with a previous TPM study [69], while here we have a vertical observation plane across all cortical layers instead of focusing on a single horizontal plane in layer 2/3. We chose 10 µA for the rest of the stimulation experiments as it was just sufficient to reliably generate an obvious neural calcium response for most sites. Note that the Thy1-GCaMP6 expression is lower in layer 4 than in other cortical layers [70] and therefore we used the dark band at around 400 µm deep as a reference marker for cortical layer 4 (Fig. 9b, c).

**Figure 9.**
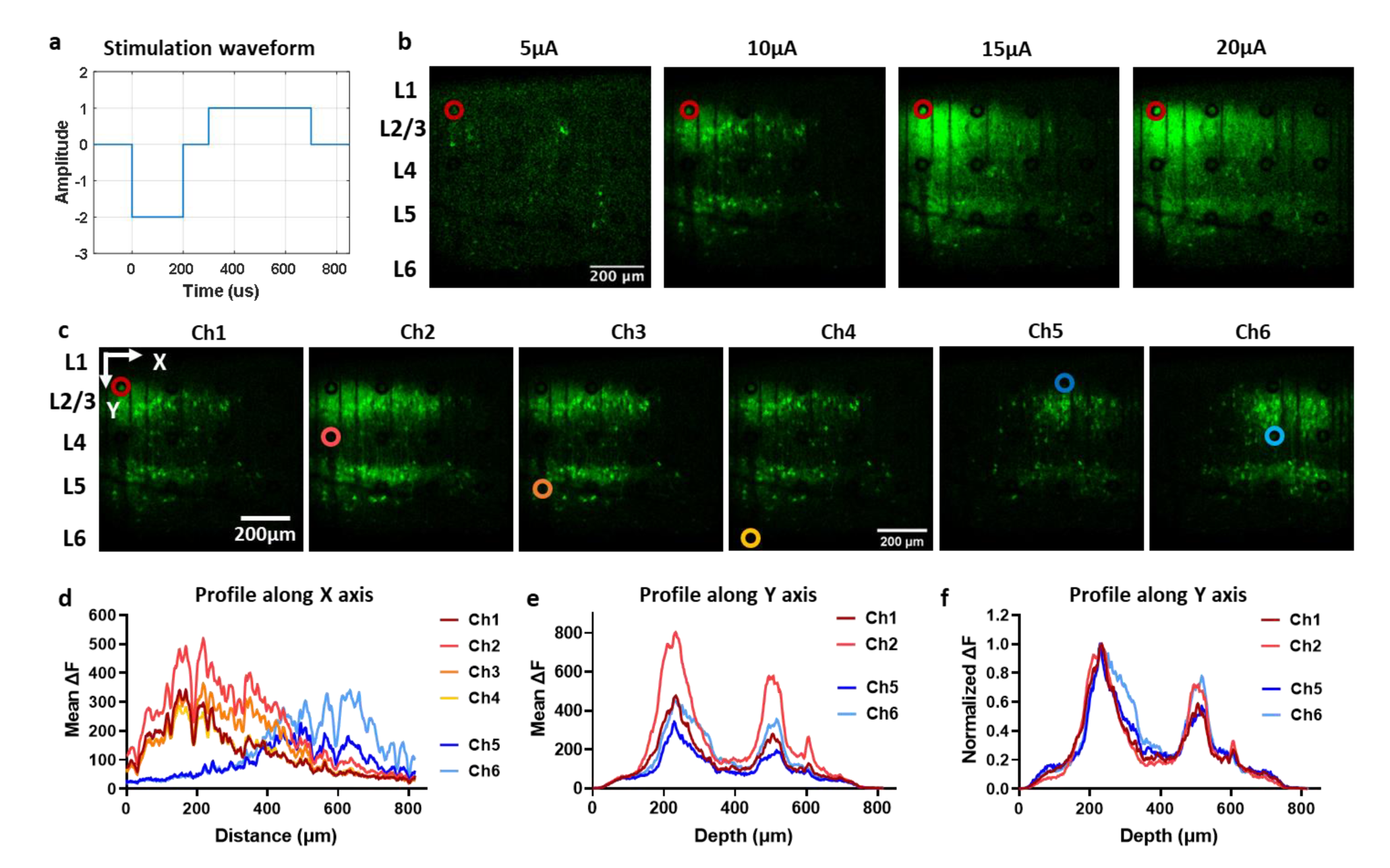
TPM imaging of the neuronal response to electrical stimulation reveals amplitude and location dependence. (a) The electrical stimulation waveform used. (b) Representative TPM images of neuronal calcium activity change in response to electrical stimulation at increasing current amplitudes. The images are averaged over three repetitions of stimulation for each amplitude. The red circle indicates the location of the stimulated channel. (c) Representative TPM images of neuronal calcium activity change in response to electrical stimulation at different locations. The images are averaged over three repetitions for each electrode site. The circles in gradient colors indicate the location of the stimulated channel. (d) The fluorescence intensity profile of images in (c) is projected along the x-axis. (e) The fluorescence intensity profile of images in (d) is projected along the y-axis. (f) Normalized fluorescence intensity profile along the y-axis.

Next, we investigated how the stimulation location affected the cortical neural network activity in response to electrical stimuli. Current pulse trains (1 s, 10 µA) were delivered at 50 Hz from the MEA while imaging neuronal calcium responses under TPM (Supplementary Video 1). We observed that electrical stimulation from the same column of the MEA activated a similar group of neurons but with different strengths (Fig. 9c, Ch1-Ch4). Additionally, electrical stimulation at a different column recruited a different cohort of neurons (Fig. 9c, Ch5-Ch6), indicating a columnar neural activation field. To qualitatively compare the strength and the distribution of the neural calcium activation, we further plotted the profile of fluorescence intensity change along the X and Y axes for the entire image (Fig. 9d, e). The layer 4 (Ch2) electrode was found to elicit the strongest GCaMP response and the layer 5 (Ch3) electrode the second strongest, while the layer 2/3 (Ch1) and layer 6 (Ch4) electrodes were lower. These results indicate that electrical stimulation may be most efficient at activating neuronal populations in general if delivered at layer 4, followed by layer 5, and less efficiently at layer 2/3 and layer 6 in this stimulation paradigm. After we normalized the fluorescence intensity to the peak intensity value for each site in Fig. 9e, we noticed higher relative peaks at around 500µm deep for Ch2 and Ch6 compared to their layer2/3 neighboring electrodes Ch1 and Ch5 (Fig. 9f). These observations indicate that apart from the overall stimulation power, layer 4 electrode sites recruit relatively more deep-layer (L5) neurons compared to layer 2/3 electrode sites. The impedance spectra of these electrode sites were similar but not completely overlapping (Supplementary Figure 1), confirming that these electrode sites were not shorted and that the difference in neural activation was unlikely to result from the variability of individual electrode sites.

In addition to the current amplitude and electrode location, frequency is also a crucial factor that can modulate the neural response to electrical stimulation. Changing the stimulation frequency has been shown to alter the type of tactile sensation and the perceived intensity in the somatosensory cortex [5, 71, 72]. There is also behavioral evidence indicating that frequency modulation of intracortical stimulation is layer-specific [73]. However, the mechanism of frequency modulation on the neural network level remains to be elucidated. With this in mind, we stimulated all the available channels of the MEA at various frequencies, ranging from 5 Hz to 150 Hz, while keeping the same amplitude (10 µA). Again, using in vivo TPM imaging through our device, we found that increasing the frequency of stimulation affects the neuronal activation pattern in a complex manner. Taking a layer 4 electrode site as an example: at 5 Hz, the neural activation was sparse and distributed; as we increased the frequency to 50 Hz, the neuronal response was much stronger, but still distributed; when we further increased the frequency, the intensity decreased, and the field of neuronal activation shrunk (Fig. 10a and b). Interestingly, although the overall neuronal activation decreased from 50 Hz to 150 Hz, more neurons at around 400 µm deep were activated at higher frequencies (Figures 10a and c). Moreover, the frequency modulation curve may also be depth dependent. Within the same column of electrodes on the MEA, the middle layer electrode sites (Ch2 and Ch3) induced the strongest overall neuronal response at ∼50 Hz, while electrodes at the superficial layer (Ch1) or the deep layer (Ch4) induced larger neuronal responses at 20 Hz compared to other frequencies tested (Fig. 10d).

**Figure 10.**
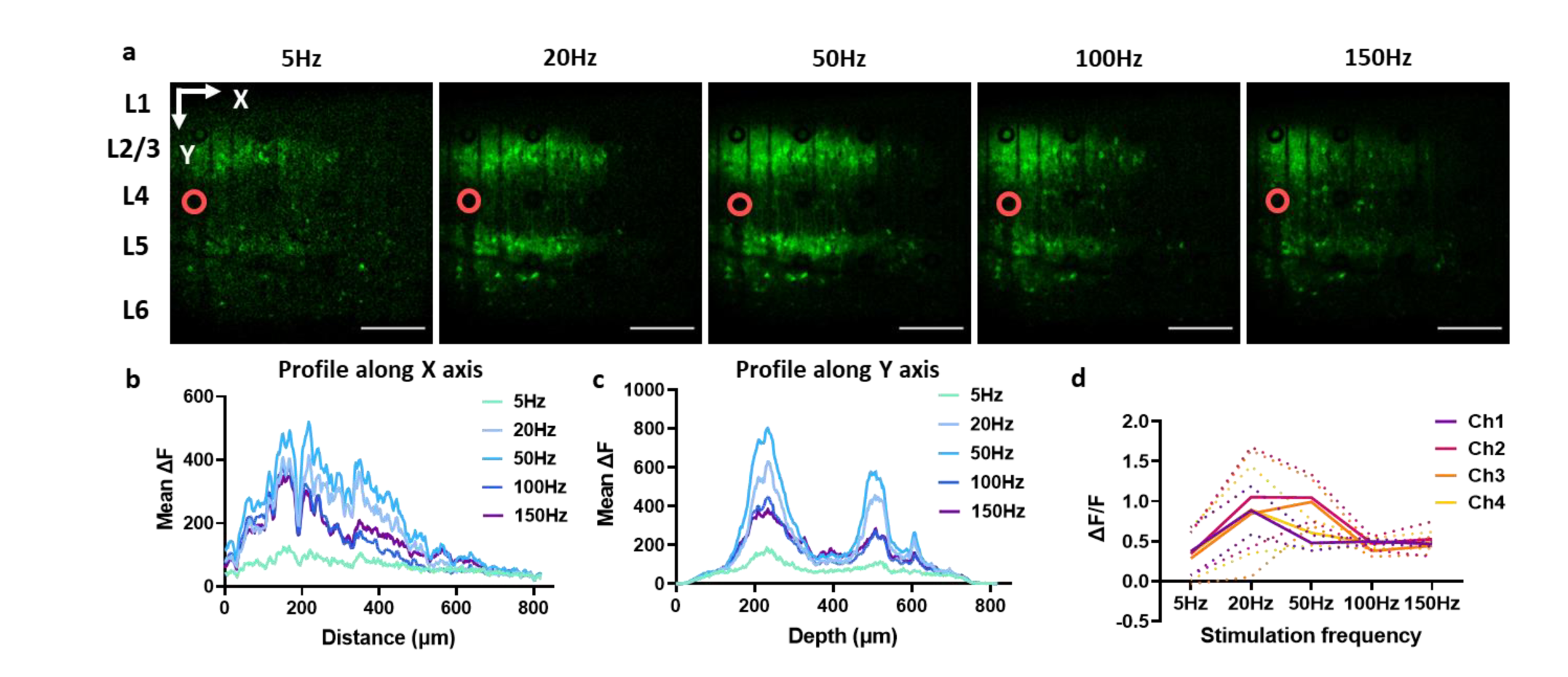
TPM imaging of the neuronal response to electrical stimulation reveals frequency dependence and depth dependence. (a) Representative TPM images of neuronal calcium activity in response to electrical stimulation from an L4 site at increasing frequencies. The stimulation consisted of bi-phasic, asymmetric, charge balanced square wave pulses at 10 µA (cathodic) for 1 sec. The images are averaged from three repetitions of stimulation. The red circle indicates the location of the stimulated channel. This imaging plane is focused at 50 µm from the MEA. Scale bar = 200 µm. (b) The fluorescence intensity profile of images in (a) is projected to the x-axis. (c) The fluorescence intensity profile of images in (a) is projected to the y-axis. (d) Frequency dependence curves of electrodes at different depths (Ch1-4). The location of Ch1-4 is the same as figure 9c. N=6 stimulations on two separate days. Data are presented as mean ± SD.

These stimulation experiments concerning depth dependence have been replicated in the same animal on different days. Although three of four animals survived at least 16 weeks of implantation (75% survival rate), two of them were excluded from the stimulation analysis due to poor imaging clarity or a liquid gap between the MEA and the brain tissue. Future studies may seek strategies to improve the biocompatibility of MEA materials and further investigate the depth dependence of intracortical microstimulation with a greater number of animals, finer step size in depth, and fluorescent labeling of other molecules of interest.

## 3. Conclusion

In this work, we developed a platform that combined a transparent MEA with a microprism to enable simultaneous electrophysiology and optical imaging in a vertical plane spanning from the brain surface to cortical layers 5/6. The custom-fabricated MEA achieves high flexibility and transparency by photolithographically patterning platinum on a flexible and transparent SU-8 substrate. In addition, the ring-shaped electrode sites and thin traces minimize the shadow of metal in the region of imaging, while reducing the photoelectrical artifact in electrophysiological recording. This novel MEA-on-microprism design remarkably expands the vertical depth of chronic two-photon imaging of electrode arrays from ∼150 µm to ∼1000 µm, visualizing multiple cortical layers simultaneously. With this setup, we were able to perform chronic TPM imaging and simultaneous electrical recording/stimulation in Thy1-GCaMP6s mice for over 4 months. High-resolution TPM images of neuronal calcium activity near the MEA have been captured at the resting state and upon sensory stimulation of the animal. We observed stable in vivo electrochemical properties of the MEA and demonstrated its capability in recording single unit activity. Thanks to the reduced photoelectric artifact, we were able to analyze the data obtained from simultaneous TPM imaging and electrophysiology recordings and correlate the calcium activity to electrical activity. Furthermore, using this device, we explored how electrode depth, stimulation amplitude, and stimulation frequency influence the neuronal calcium responses across all cortical layers to intracortical microstimulation. To the best of our knowledge, this is the first in vivo TPM imaging work that directly visualizes the cortical multi-layer neuronal network in response to microstimulation. Our work provides a facile but powerful design for concurrent electrophysiology and optical imaging throughout all cortical layers, which could revolutionize the study of cellular mechanisms across large-scale cortical networks in neuroscience research and neural prosthetic device development.

## 4. Methods

### 4.1. Fabrication of MEA

Microelectrode array (MEA) fabrication: A Si wafer with a 1µm thick SiO_2_ layer (University Wafer Inc) was first cleaned by sonicating in acetone, isopropanol, and DI water sequentially for 5 mins at each step. The wafer was then dried on a hot plate at 150 µC for 3 mins and cleaned by O_2_ plasma using a reactive ion etcher (RIE, Trion Phantom III LT) for 120 s at 200 mTorr pressure and 150 Watts power. The cleaned wafer was spin-coated with SU-8 2015 (MicroChemicals) at 5000 rpm for 1 min and soft baked at 65°C for 3 mins and 95°C for 5 mins. Then the wafer was exposed using a maskless aligner (MLA, MLA100 Heidelberg Instruments) with a dose of 400 mJ/cm^2^. After exposure, the SU-8 first layer was post-baked at 65°C for 3 mins and 95°C for 5 mins, developed using SU-8 developer (MicroChemicals) for 1 min and cleaned by isopropanol and DI water, and hard baked at 200 µC, 180 µC, and 150°C for 5 mins each and allowed to cool down below 95°C. The wafer was then treated with O_2_ plasma with RIE for 75 s at a pressure of 200 mTorr and 150W power. The cleaned wafer was then spin-coated with AZ P4620 photoresist (MicroChemicals) at 5300 rpm for 1min and baked at 105°C for 5 mins. After baking the wafer is exposed using MLA with a dose of 700 mJ/cm^2^, then developed using AZ400k 1:4 developer (MicroChemicals), cleaned by water rinse, and dried by N_2_ gas flow. A mild 60s RIE O_2_ plasma cleaning at pressure 600 mTorr and 60 W power was performed before metal deposition. 10 nm Ti adhesion layer and 100 nm Pt layer were evaporated on the wafer using an Electron Beam Evaporator (Plassys MEB550S). Then the metal was lifted-off in acetone overnight. The next day, the wafer was cleaned by O_2_ plasma for 60 s at 600 mTorr and 60 W, then spin-coated with SU-8 2015 at 5000 rpm for 1 min and soft baked at 65°C for 2 mins and 95°C for 5 mins. The wafer was exposed using MLA with a dose of 400 mJ/cm^2^, post-baked, and developed with an SU-8 developer. The wafer was then cleaned with isopropanol and water, and hard baked at 200 µC, 180 µC, and 150°C for 5 mins each and allowed to cool down below 95°C. The MEAs were lifted off from the wafer using buffered oxide etchant (1:7) in an acid hood for 8 hours.

### 4.2. Electrodeposition of Pt

Nano-platinum was electrochemically formed starting from a solution containing 25 mM H_2_(PtCl_6_) in 1 mM HCl, by applying a constant voltage at −0.3 V for 200 s. Electrochemical depositions were carried out using a potentiostat/galvanostat (Autolab, Metrohm, USA), connected to a three-electrode electrochemical cell with a platinum counter electrode and an Ag/AgCl reference electrode.

### 4.3. Assembly of microprism and MEA

Similar to previous work, microprisms were purchased (#MPCH-1.0, Tower Optical Corporation) and adhered to three layers of cover glasses (two 3×3 mm square, one 5 mm round, #1 thickness, 0.15 ± 0.02 mm, Warner Instruments LLC) using a transparent glue (NOA 71, Norland Optical).

The printed circuit board (PCB) was manufactured following a customized design. The MEA was connected to the PCB using a silver heat-seal connector (P/N HST-9805-210, Elform). Specifically, the connector cable was trimmed to fit the width of PCB and MEA with a length of 1∼2cm. The cable was then aligned and bonded to the PCB and MEA using a hair straightener heated to 380 ^µ^F for ∼30s with pressure. Two pieces of PDMS (∼1mm thick) were used to pad between the heating pad and the PCB/MEA on both sides. After electrochemically depositing nano platinum onto the sites, we applied a drop of NOA 71 glue to the back side of the MEA tip and carefully adhered the MEA to the microprism imaging face and the cover glasses, with an angled tip tweezer holding the MEA bent at ∼90 degrees at the corner. Excess glue that ran to the front side of MEA was gently removed with a fine tip tweezer. The ground and reference site on the PCB was connected to a silver wire, to be wired to the bone screw on the animal later. Finally, all the exposed traces and fragile parts were covered with a thin layer of silicone to provide additional insulation and mechanical support. Fully assembled devices were characterized with electrochemical methods described below and sterilized with the ethylene oxide sterilization process before surgery.

### 4.4. Electrochemical characterizations

Electrochemical impedance spectroscopy (EIS) and cyclic voltammetry (CV) measurements were used to investigate the electrode/solution interface in vitro, before implantation, and in vivo.

During the EIS measurements, a sine wave (10 mV RMS amplitude) was superimposed onto the open circuit potential while varying the frequency from 1 to 10^5^ Hz. During the CV tests, the working electrode potential was swept between 0.8V and −0.6 V with a scan rate of 1 V/s. Both EIS and CV measurements were carried out using a potentiostat/galvanostat (Autolab, Metrohm, USA).

In vitro, EIS and CV were performed in 1x phosphate buffered saline (PBS, composition: 11.9 mM Na_2_HPO_4_ and KH_2_PO_4_, 137 mM NaCl, 2.7 mM KCl, pH 7.4) using a three-electrode electrochemical cell configuration with a platinum counter electrode and an Ag/AgCl reference electrode.

In the brain, EIS and CV were measured using a two-electrode electrochemical cell configuration, where a stainless-steel bone screw, anchored on the contralateral skull, served as the counter and reference electrode.

### 4.5. Animals and surgery

Transgenic mice expressing the neuronal activity reporter GCaMP6s in cortical pyramidal neurons (male, n=3, strain Tg (Thy1-GCaMP6s) GP4.3Dkim) were used in this study and obtained from Jackson Laboratories (Bar Harbor, ME USA). During surgery, mice were anesthetized using ketamine (75 mg/kg)/xylazine(10 mg/kg) and maintained with updates of ketamine (22.5 mg/kg) alone as needed. Animal fur on top of the head was trimmed off and the skin surface was then sterilized with 70% isopropyl alcohol and betadine. The animal head was fixed onto a stereotaxic frame (Narishige International USA) and the temperature was kept at 37 °C with an electric heating pad. After skin resection and skull drying with Vetbond (3M, St. Paul, MN), a rectangular stainless-steel frame (#CF-10, Narishige International USA) was adhered to the skull with dental cement (A-M Systems, Sequim, WA). We waited until the dental cement completely solidified before transferring the animal to a stereotaxic frame-holder (Narishige International USA). A stainless-steel bone screw (0.86 mm shaft diameter, Fine Science Tools, Inc.) was secured on the left parietal bone of the skull as reference and ground for the electrode array. Using a high-speed dental drill, we created a 4×4 mm craniotomy above the somatosensory and visual cortex over the right hemisphere. The dura mater within the cranial window was removed with a fine tip tweezer and a bent 30-gauge needle. A lance-shaped 0.1 mm thick razor blade (#72000, Electron Microscopy Sciences) was attached to a stereotaxic arm and inserted vertically, then moved laterally to create a 1×1 mm incision. We applied saline and Gelfoam to the surface of the brain and waited for the bleeding to stop. The PCB of the device was anchored to a stereotaxic arm angled at about 30 degrees approaching the cranial window from the back of the animal. Another stereotaxic arm was oriented vertically to hold the cover glass of the microprism/MEA with a vacuum line. The microprism/MEA was inserted vertically into the incision, with the imaging surface of the microprism facing the posterior (visual cortex) of the animal. The microprism/MEA device was lowered until the round coverglass touched the skull so that the square cover glass was contacting the brain surface with a little pressure to prevent dura and meningeal regrowth. Tiny drops of silicone sealant (Kwik-sil, World Precision Instruments) were applied around the coverglass and on the uncovered brain surface, if any, to seal the cranial window. The vacuum was released after dental cement secured the implant in place. The ground/reference wire was connected to the bone screw. The excess cable was folded to the back of the PCB and covered with dental cement. After surgery, the animal was treated with 5 mg/kg ketoprofen (100 mg/ ml, Zoetis Inc., Kalamazoo, MI) and 10 mg/kg Baytril solution (Henry Schein Inc.) for three days. All experimental protocols were approved by the University of Pittsburgh, Division of Laboratory Animal Resources and Institutional Animal Care and Use Committee (ARO ID: IS00018691) in accordance with the standards for humane animal care as set by the Animal Welfare Act and the National Institutes of Health Guide for the Care and Use of Laboratory Animals.

### 4.6. In vivo two-photon imaging and image processing

The animal was head-fixed, free-running on a custom treadmill during two-photon imaging. The two-photon imaging system has been described in our previous publications [27, 53]. It consists of an ultra-fast laser (Insight DS+; Spectra-Physics, Menlo Park, CA), a scan head (Bruker, Madison, WI), non-descanned photomultiplier tubes (Hamamatsu Photonics KK, Hamamatsu, Shizuoka, Japan), and a 16X 0.8 NA water immersion objective lens (Nikon Instruments, Melville, NY). The excitation laser was set at a wavelength of 920 nm. Sulforhodamine 101 (SR101) (∼0.05 cc; 1 mg/ml) was injected before imaging for visualization of blood vessels. For imaging neuronal calcium activity, time-series images were collected at 512 x 512 x 1 pixel and at ∼1 frame per second (fps).

The time-series images were processed and analyzed using ImageJ and customized IJ1 macro scripts. The images were first denoised with a median filter and the “Despeckle” function. A background image was acquired by averaging the whole image stack or the images before any stimulation. Then we subtracted the background image from every image in the time series. The resulting fluorescence change images were maximum intensity projected to a single image to capture all active neurons during the period of interest. The vasculature in Fig. 4 is the average image of the red (SR101) channel, which is superimposed on the above neural activity image in the green (Thy1-GCaMP) channel.

For quantification of neuronal calcium intensity change in Figure 8, we manually circled the neuronal soma from the raw image stack and exported the intensity change over time. The fluorescent intensity change ratio (ΔF/F) was then calculated, where F was the mean value of fluorescence intensity in the first 30s before light stimulation, and ΔF was the difference between F and the fluorescence at each time step. In Fig. 9 and Fig. 10, the fluorescent intensity profile of the entire image along the x and y axis is measured with the “Plot profile” function in ImageJ. The image was rotated 90 degrees to change the direction of the profile projection.

### 4.7. Electrophysiological recording

The electrophysiological recording was performed on awake animals. Data was collected using a Ripple Grapevine (Nano2+stim Ripple LLC, Salt Lake City, Utah) and Trellis software at a 30 kHz sample rate. Spike waveforms were sorted in Plexon Offline Sorter (version 3) using the spike train data. We manually sorted single units from 3-D PCA clusters. Noise waveforms that co-occurred in multiple channels were excluded.

### 4.8. Electrical stimulation

Current-controlled stimulation was delivered using an Autolab (Metrohm, USA). 1-sec long pulse trains containing biphasic, charge-balanced asymmetric square waves with various amplitudes and frequencies were delivered. Each pulse had a 200 µs cathodic phase, a 100 µs interval, and a 400 µs anodic phase. The anodic current amplitude was set at half of the cathodic current. During one imaging session, each electrode site was stimulated three times in a row; each time there was a 1-sec stimulation period and a 3-sec off period (25% duty cycle).

## 5 Acknowledgement

The authors thank Dr. Hui Fang and Dr. Qiang Yi for advice and materials on the heat-seal connector. The authors thank Daniela Krahe for occasional help with recordings. This work was supported by NIH R01NS089688 and BRAIN R01NS110564.

## 6. Table of Contents

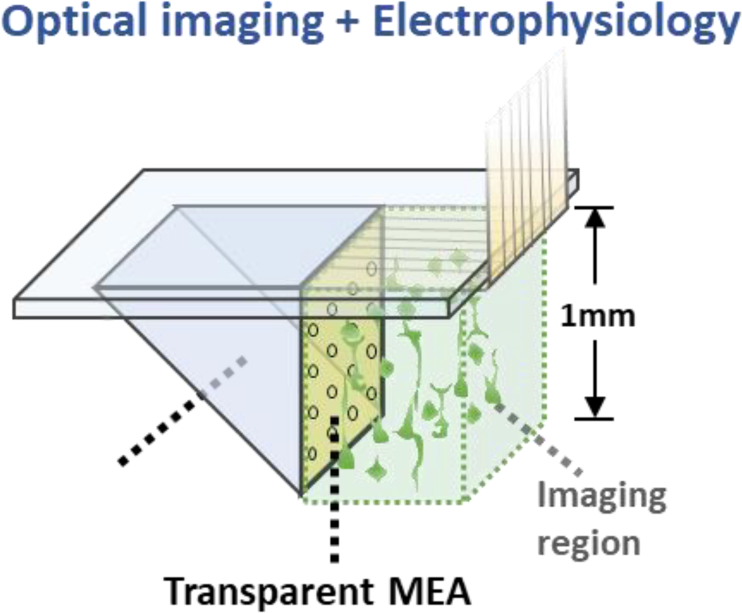

This MEA-on-microprism design integrates a 16-channel transparent electrode array with a microprism to allow simultaneous electrophysiology and optical imaging throughout all cortical layers. By implanting this device in the visual cortex of mice, we achieved single-unit recording, microstimulation, and two-photon imaging of the whole cortical column for over 4 months. Concurrent microstimulation and two-photon imaging revealed depth dependency of various stimulation parameters.

## Supplementary materials

**Supplementary table 1.** Comparison of the size and impedance of transparent MEAs.

| | Electrode area<br>($\mu\text{m}^2$) | Impedance at 1kHz<br>($\text{k}\Omega$) | Area specific<br>impedance ($\Omega \cdot \text{cm}^2$) |
| --- | --- | --- | --- |
| Ring Pt (this work) | 706.86 | 41.34 | 0.292215924 |
| Au/PEDOT:PSS<br>nanomesh [43] | 1256.64 | 43.4 | 0.54538176 |
| ITO [40] | 49087 | 20 | 9.8174 |
| Doped graphene [35] | 2500 | 541 | 13.525 |
| 4-layer graphene [36] | 7854 | 286.4 | 22.493856 |
| PtNP coated graphene<br>[60] | 10000 | 872.9 | 87.29 |
| Graphene [37] | 10000 | 963 | 96.3 |

**Supplementary figure 1.**
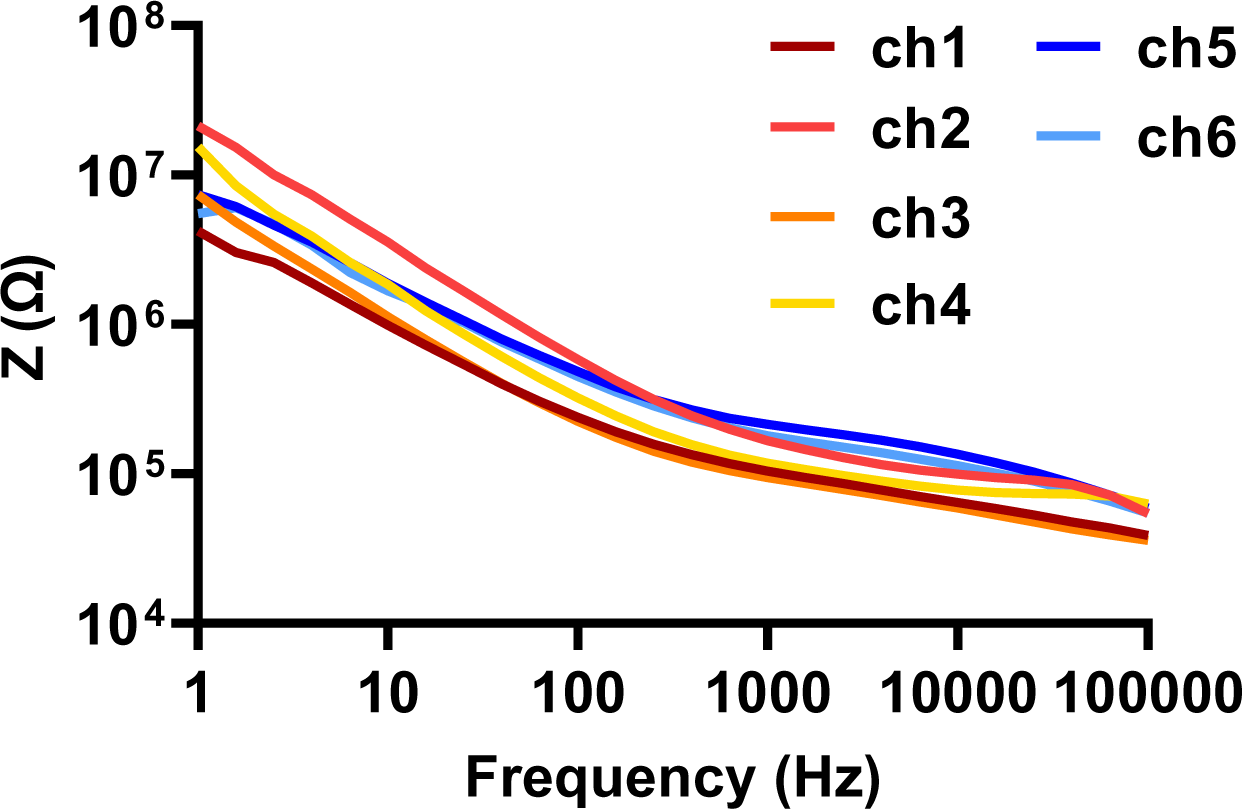
Impedance spectrum of the electrode sites presented in figure 8 on the day of stimulation. Supplementary materials

## References

[1] S.M. Schuetze, The discovery of the action potential, Trends in Neurosciences 6 (1983) 164-168.

[2] H. Asanuma, S. Stoney Jr, C. Abzug, Relationship between afferent input and motor outflow in cat motorsensory cortex, Journal of Neurophysiology 31(5) (1968) 670-681.

[3] W.G. Penfield, Ferrier lecture-some observations on the cerebral cortex of man, Proceedings of the Royal Society of London. Series B-Biological Sciences 134(876) (1947) 329-347.

[4] E. Bizzi, Discharge of frontal eye field neurons during saccadic and following eye movements in unanesthetized monkeys, Experimental Brain Research 6(1) (1968) 69-80.

[5] S.N. Flesher, J.L. Collinger, S.T. Foldes, J.M. Weiss, J.E. Downey, E.C. Tyler-Kabara, S.J. Bensmaia, A.B. Schwartz, M.L. Boninger, R.A. Gaunt, Intracortical microstimulation of human somatosensory cortex, Science translational medicine 8(361) (2016) 361ra141-361ra141.

[6] G.A. Tabot, J.F. Dammann, J.A. Berg, F.V. Tenore, J.L. Boback, R.J. Vogelstein, S.J. Bensmaia, Restoring the sense of touch with a prosthetic hand through a brain interface, Proceedings of the National Academy of Sciences 110(45) (2013) 18279-18284.

[7] W.H. Dobelle, Artificial vision for the blind by connecting a television camera to the visual cortex, ASAIO journal 46(1) (2000) 3-9.

[8] J.L. Collinger, B. Wodlinger, J.E. Downey, W. Wang, E.C. Tyler-Kabara, D.J. Weber, A.J. McMorland, M. Velliste, M.L. Boninger, A.B. Schwartz, High-performance neuroprosthetic control by an individual with tetraplegia, The Lancet 381(9866) (2013) 557-564.

[9] D.M. Taylor, S.I.H. Tillery, A.B. Schwartz, Direct cortical control of 3D neuroprosthetic devices, Science 296(5574) (2002) 1829-1832.

[10] G. Buzsáki, Large-scale recording of neuronal ensembles, Nature neuroscience 7(5) (2004) 446-451.

[11] W. Denk, J.H. Strickler, W.W. Webb, Two-photon laser scanning fluorescence microscopy, Science 248(4951) (1990) 73-76.

[12] K. Svoboda, W. Denk, D. Kleinfeld, D.W. Tank, In vivo dendritic calcium dynamics in neocortical pyramidal neurons, Nature 385(6612) (1997) 161-165.

[13] C. Stosiek, O. Garaschuk, K. Holthoff, A. Konnerth, In vivo two-photon calcium imaging of neuronal networks, Proceedings of the National Academy of Sciences 100(12) (2003) 7319-7324.

[14] A. Song, A.S. Charles, S.A. Koay, J.L. Gauthier, S.Y. Thiberge, J.W. Pillow, D.W. Tank, Volumetric two-photon imaging of neurons using stereoscopy (vTwINS), Nature methods 14(4) (2017) 420-426.

[15] M.B. Bouchard, V. Voleti, C.S. Mendes, C. Lacefield, W.B. Grueber, R.S. Mann, R.M. Bruno, E.M. Hillman, Swept confocally-aligned planar excitation (SCAPE) microscopy for high-speed volumetric imaging of behaving organisms, Nature photonics 9(2) (2015) 113-119.

[16] W. Zong, H.A. Obenhaus, E.R. Skytøen, H. Eneqvist, N.L. de Jong, R. Vale, M.R. Jorge, M.-B. Moser, E.I. Moser, Large-scale two-photon calcium imaging in freely moving mice, Cell 185(7) (2022) 1240-1256. e30.

[17] L. Ghanbari, R.E. Carter, M.L. Rynes, J. Dominguez, G. Chen, A. Naik, J. Hu, M.A.K. Sagar, L. Haltom, N. Mossazghi, Cortex-wide neural interfacing via transparent polymer skulls, Nature communications 10(1) (2019) 1-13.

[18] A.A. Oliva, M. Jiang, T. Lam, K.L. Smith, J.W. Swann, Novel hippocampal interneuronal subtypes identified using transgenic mice that express green fluorescent protein in GABAergic interneurons, Journal of Neuroscience 20(9) (2000) 3354-3368.

[19] Y. Wang, T. Kakizaki, H. Sakagami, K. Saito, S. Ebihara, M. Kato, M. Hirabayashi, Y. Saito, N. Furuya, Y. Yanagawa, Fluorescent labeling of both GABAergic and glycinergic neurons in vesicular GABA transporter (VGAT)–Venus transgenic mouse, Neuroscience 164(3) (2009) 1031-1043.

[20] N.C. Flytzanis, C.N. Bedbrook, H. Chiu, M.K. Engqvist, C. Xiao, K.Y. Chan, P.W. Sternberg, F.H. Arnold, V. Gradinaru, Archaerhodopsin variants with enhanced voltage-sensitive fluorescence in mammalian and Caenorhabditis elegans neurons, Nature communications 5(1) (2014) 1-9.

[21] A. Nimmerjahn, F. Helmchen, In vivo labeling of cortical astrocytes with sulforhodamine 101 (SR101), Cold Spring Harbor Protocols 2012(3) (2012) pdb. prot068155.

[22] I. Ferezou, S. Bolea, C.C. Petersen, Visualizing the cortical representation of whisker touch: voltage-sensitive dye imaging in freely moving mice, Neuron 50(4) (2006) 617-629.

[23] H. Wang, M. Jing, Y. Li, Lighting up the brain: genetically encoded fluorescent sensors for imaging neurotransmitters and neuromodulators, Current Opinion in Neurobiology 50 (2018) 171-178.

[24] G. Miesenböck, D.A. De Angelis, J.E. Rothman, Visualizing secretion and synaptic transmission with pH-sensitive green fluorescent proteins, Nature 394(6689) (1998) 192-195.

[25] S. Okumoto, L.L. Looger, K.D. Micheva, R.J. Reimer, S.J. Smith, W.B. Frommer, Detection of glutamate release from neurons by genetically encoded surface-displayed FRET nanosensors, Proceedings of the National Academy of Sciences 102(24) (2005) 8740-8745.

[26] J.R. Eles, A.L. Vazquez, T.D. Kozai, X.T. Cui, In vivo imaging of neuronal calcium during electrode implantation: spatial and temporal mapping of damage and recovery, Biomaterials 174 (2018) 79-94.

[27] T.D.Y. Kozai, A.L. Vazquez, C.L. Weaver, S.-G. Kim, X.T. Cui, In vivo two-photon microscopy reveals immediate microglial reaction to implantation of microelectrode through extension of processes, Journal of neural engineering 9(6) (2012) 066001.

[28] X.S. Zheng, Q. Yang, A. Vazquez, X.T. Cui, Imaging the efficiency of PEDOT/CNT and iridium oxide electrode coatings for microstimulation.

[29] K.C. Stieger, J.R. Eles, K.A. Ludwig, T.D. Kozai, In vivo microstimulation with cathodic and anodic asymmetric waveforms modulates spatiotemporal calcium dynamics in cortical neuropil and pyramidal neurons of male mice, Journal of neuroscience research 98(10) (2020) 2072-2095.

[30] N.J. Michelson, J.R. Eles, A.L. Vazquez, K.A. Ludwig, T.D. Kozai, Calcium activation of cortical neurons by continuous electrical stimulation: Frequency dependence, temporal fidelity, and activation density, Journal of neuroscience research 97(5) (2019) 620-638.

[31] J.R. Eles, T.D. Kozai, In vivo imaging of calcium and glutamate responses to intracortical microstimulation reveals distinct temporal responses of the neuropil and somatic compartments in layer II/III neurons, Biomaterials 234 (2020) 119767.

[32] T.D. Kozai, J.R. Eles, A.L. Vazquez, X.T. Cui, Two-photon imaging of chronically implanted neural electrodes: Sealing methods and new insights, Journal of neuroscience methods 258 (2016) 46-55.

[33] J. Eles, A. Vazquez, T. Kozai, X. Cui, Meningeal inflammatory response and fibrous tissue remodeling around intracortical implants: an in vivo two-photon imaging study, Biomaterials 195 (2019) 111-123.

[34] P. Ledochowitsch, A. Yazdan-Shahmorad, K. Bouchard, C. Diaz-Botia, T. Hanson, J.-W. He, B. Seybold, E. Olivero, E. Phillips, T. Blanche, Strategies for optical control and simultaneous electrical readout of extended cortical circuits, Journal of neuroscience methods 256 (2015) 220-231.

[35] D. Kuzum, H. Takano, E. Shim, J.C. Reed, H. Juul, A.G. Richardson, J. De Vries, H. Bink, M.A. Dichter, T.H. Lucas, Transparent and flexible low noise graphene electrodes for simultaneous electrophysiology and neuroimaging, Nature communications 5(1) (2014) 1-10.

[36] D.-W. Park, J.P. Ness, S.K. Brodnick, C. Esquibel, J. Novello, F. Atry, D.-H. Baek, H. Kim, J. Bong, K.I. Swanson, Electrical neural stimulation and simultaneous in vivo monitoring with transparent graphene electrode arrays implanted in GCaMP6f mice, ACS nano 12(1) (2018) 148-157.

[37] M. Thunemann, Y. Lu, X. Liu, K. Kılıç, M. Desjardins, M. Vandenberghe, S. Sadegh, P.A. Saisan, Q. Cheng, K.L. Weldy, Deep 2-photon imaging and artifact-free optogenetics through transparent graphene microelectrode arrays, Nature communications 9(1) (2018) 1-12.

[38] D.-W. Park, A.A. Schendel, S. Mikael, S.K. Brodnick, T.J. Richner, J.P. Ness, M.R. Hayat, F. Atry, S.T. Frye, R. Pashaie, Graphene-based carbon-layered electrode array technology for neural imaging and optogenetic applications, Nature communications 5(1) (2014) 1-11.

[39] K.Y. Kwon, B. Sirowatka, A. Weber, W. Li, Opto-μECoG array: A hybrid neural interface with transparent μECoG electrode array and integrated LEDs for optogenetics, IEEE transactions on biomedical circuits and systems 7(5) (2013) 593-600.

[40] N. Kunori, I. Takashima, A transparent epidural electrode array for use in conjunction with optical imaging, Journal of neuroscience methods 251 (2015) 130-137.

[41] C.F. Guo, T. Sun, Q. Liu, Z. Suo, Z. Ren, Highly stretchable and transparent nanomesh electrodes made by grain boundary lithography, Nature communications 5(1) (2014) 1-8.

[42] W. Lee, D. Kim, N. Matsuhisa, M. Nagase, M. Sekino, G.G. Malliaras, T. Yokota, T. Someya, Transparent, conformable, active multielectrode array using organic electrochemical transistors, Proceedings of the National Academy of Sciences 114(40) (2017) 10554-10559.

[43] Y. Qiang, P. Artoni, K.J. Seo, S. Culaclii, V. Hogan, X. Zhao, Y. Zhong, X. Han, P.-M. Wang, Y.-K. Lo, Transparent arrays of bilayer-nanomesh microelectrodes for simultaneous electrophysiology and two-photon imaging in the brain, Science advances 4(9) (2018) eaat0626.

[44] C.D. Gilbert, Microcircuitry of the visual cortex, Annual review of neuroscience (1983).

[45] R.J. Douglas, K. Martin, A functional microcircuit for cat visual cortex, The Journal of physiology 440(1) (1991) 735-769.

[46] Y. Amir, M. Harel, R. Malach, Cortical hierarchy reflected in the organization of intrinsic connections in macaque monkey visual cortex, Journal of Comparative Neurology 334(1) (1993) 19-46.

[47] T. Binzegger, R.J. Douglas, K.A. Martin, A quantitative map of the circuit of cat primary visual cortex, Journal of Neuroscience 24(39) (2004) 8441-8453.

[48] J.M. Blackwell, M.N. Geffen, Progress and challenges for understanding the function of cortical microcircuits in auditory processing, Nature communications 8(1) (2017) 1-9.

[49] T.H. Chia, M.J. Levene, Microprisms for in vivo multilayer cortical imaging, Journal of neurophysiology 102(2) (2009) 1310-1314.

[50] M.L. Andermann, N.B. Gilfoy, G.J. Goldey, R.N. Sachdev, M. Wölfel, D.A. McCormick, R.C. Reid, M.J. Levene, Chronic cellular imaging of entire cortical columns in awake mice using microprisms, Neuron 80(4) (2013) 900-913.

[51] R.J. Low, Y. Gu, D.W. Tank, Cellular resolution optical access to brain regions in fissures: imaging medial prefrontal cortex and grid cells in entorhinal cortex, Proceedings of the National Academy of Sciences 111(52) (2014) 18739-18744.

[52] L. Beckmann, X. Zhang, N.A. Nadkarni, Z. Cai, A. Batra, D.P. Sullivan, W.A. Muller, C. Sun, R. Kuranov, H.F. Zhang, Longitudinal deep-brain imaging in mouse using visible-light optical coherence tomography through chronic microprism cranial window, Biomedical optics express 10(10) (2019) 5235-5250.

[53] Q. Yang, A.L. Vazquez, X.T. Cui, Long-term in vivo two-photon imaging of the neuroinflammatory response to intracortical implants and micro-vessel disruptions in awake mice, Biomaterials 276 (2021) 121060.

[54] M. Hopcroft, T. Kramer, G. Kim, K. Takashima, Y. Higo, D. Moore, J. Brugger, Micromechanical testing of SU-8 cantilevers, Fatigue & Fracture of Engineering Materials & Structures 28(8) (2005) 735-742.

[55] S.-H. Huang, S.-P. Lin, J.-J.J. Chen, In vitro and in vivo characterization of SU-8 flexible neuroprobe: From mechanical properties to electrophysiological recording, Sensors and Actuators A: Physical 216 (2014) 257-265.

[56] Z. Chen, J.-B. Lee, Biocompatibility of su-8 and its biomedical device applications, Micromachines 12(7) (2021) 794.

[57] G.S. Brindley, W.S. Lewin, The sensations produced by electrical stimulation of the visual cortex, The Journal of physiology 196(2) (1968) 479-493.

[58] S.B. Brummer, M. Turner, Electrochemical considerations for safe electrical stimulation of the nervous system with platinum electrodes, IEEE Transactions on Biomedical Engineering (1) (1977) 59-63.

[59] A. Merz, A.J. Bard, A stable surface modified platinum electrode prepared by coating with electroactive polymer, Journal of the American Chemical Society 100(10) (1978) 3222-3223.

[60] Y. Lu, X. Liu, R. Hattori, C. Ren, X. Zhang, T. Komiyama, D. Kuzum, Ultralow impedance graphene microelectrodes with high optical transparency for simultaneous deep two-photon imaging in transgenic mice, Advanced functional materials 28(31) (2018) 1800002.

[61] T.D. Kozai, A.L. Vazquez, Photoelectric artefact from optogenetics and imaging on microelectrodes and bioelectronics: new challenges and opportunities, Journal of Materials Chemistry B 3(25) (2015) 4965-4978.

[62] L.A. Palmer, T.L. Davis, Receptive-field structure in cat striate cortex, Journal of neurophysiology 46(2) (1981) 260-276.

[63] E.D. Adrian, R. Matthews, The action of light on the eye: Part I. The discharge of impulses in the optic nerve and its relation to the electric changes in the retina, The Journal of Physiology 63(4) (1927) 378.

[64] W. Bair, J.R. Cavanaugh, M.A. Smith, J.A. Movshon, The timing of response onset and offset in macaque visual neurons, Journal of Neuroscience 22(8) (2002) 3189-3205.

[65] S.R. Olsen, D.S. Bortone, H. Adesnik, M. Scanziani, Gain control by layer six in cortical circuits of vision, Nature 483(7387) (2012) 47-52.

[66] G. DeAngelis, J. Robson, I. Ohzawa, R. Freeman, Organization of suppression in receptive fields of neurons in cat visual cortex, Journal of Neurophysiology 68(1) (1992) 144-163.

[67] D.S. Bortone, S.R. Olsen, M. Scanziani, Translaminar inhibitory cells recruited by layer 6 corticothalamic neurons suppress visual cortex, Neuron 82(2) (2014) 474-485.

[68] E.J. Tehovnik, Electrical stimulation of neural tissue to evoke behavioral responses, Journal of neuroscience methods 65(1) (1996) 1-17.

[69] M.H. Histed, V. Bonin, R.C. Reid, Direct activation of sparse, distributed populations of cortical neurons by electrical microstimulation, Neuron 63(4) (2009) 508-522.

[70] H. Dana, T.-W. Chen, A. Hu, B.C. Shields, C. Guo, L.L. Looger, D.S. Kim, K. Svoboda, Thy1-GCaMP6 transgenic mice for neuronal population imaging in vivo, PloS one 9(9) (2014) e108697.

[71] C.L. Hughes, S.N. Flesher, J.M. Weiss, M. Boninger, J.L. Collinger, R.A. Gaunt, Perception of microstimulation frequency in human somatosensory cortex, ELife 10 (2021) e65128.

[72] T. Callier, N.W. Brantly, A. Caravelli, S.J. Bensmaia, The frequency of cortical microstimulation shapes artificial touch, Proceedings of the National Academy of Sciences 117(2) (2020) 1191-1200.

[73] M.E. Urdaneta, N.G. Kunigk, F. Delgado, S.I. Fried, K.J. Otto, Layer-specific parameters of intracortical microstimulation of the somatosensory cortex, Journal of Neural Engineering 18(5) (2021) 055007.

